# Changes in excitability properties of ventromedial motor thalamic neurons in 6-OHDA lesioned mice

**DOI:** 10.1101/2020.10.05.327379

**Authors:** Edyta K Bichler, Francesco Cavarretta, Dieter Jaeger

## Abstract

The activity of basal ganglia input receiving motor thalamus (BGMT) makes a critical impact on motor cortical processing, but modification in BGMT processing with Parkinsonian conditions have not be investigated at the cellular level. Such changes may well be expected due to homeostatic regulation of neural excitability in the presence of altered synaptic drive with dopamine depletion. We addressed this question by comparing BGMT properties in brain slice recordings between control and unilaterally 6-OHDA treated adult mice. At a minimum of 1 month post 6-OHDA treatment, BGMT neurons showed a highly significant increase in intrinsic excitability, which was primarily due to a decrease in M-type potassium current. BGMT neurons after 6-OHDA treatment also showed an increase in T-type calcium rebound spikes following hyperpolarizing current steps. Biophysical computer modeling of a thalamic neuron demonstrated that an increase in rebound spiking can also be accounted for by a decrease in the M-type potassium current. Modeling also showed that an increase in sag with hyperpolarizing steps found after 6-OHDA treatment could in part but not fully be accounted for by the decrease in M-type current. These findings support the hypothesis that homeostatic changes in BGMT neural properties following 6-OHDA treatment likely influence the signal processing taking place in basal ganglia thalamocortical processing in Parkinson’s disease.

**Significance Statement:** Our investigation of the excitability properties of neurons in the basal ganglia input receiving motor thalamus (BGMT) is significant because they are likely to be different from properties in other thalamic nuclei due to the additional inhibitory input stream these neurons receive. Further, they are important to understand the role of BGMT in the dynamic dysfunction of cortico – basal ganglia circuits in Parkinson’s disease. We provide clear evidence that after 6-OHDA treatment of mice important homeostatic changes occur in the intrinsic properties of BGMT neurons. Specifically we identify the M-type potassium current as an important thalamic excitability regulator in the parkinsonian state.

## Introduction

The basal ganglia (BG) form strong connections with motor and premotor cerebral cortical areas through output from the Substantia Nigra pars reticulata (SNr) and internal globus pallidus (GPi) (Alexander and Crutcher, 1990; Alexander et al., 1986). These BG outputs terminate as GABAergic inhibitory connections in the motor thalamus, in rodents primarily in the ventromedial and ventroanterior nuclei (VM/VAL) (Bosch-Bouju et al., 2013; Kuramoto et al., 2011; Kuramoto et al., 2009). We refer to this as the BG input recipient motor thalamus (BGMT). Glutamatergic thalamocortical neurons in BGMT project to motor and premotor cortex, where they connect primarily to pyramidal neuron dendrites in layer 1 (Guo et al., 2018; Kuramoto et al., 2009; Kuramoto et al., 2015). In traditional models of BG function, the motor thalamus acts purely as a relay, and transmits a spike rate code by which excessive movement is suppressed through tonic BG inhibition of motor thalamus (Alexander and Crutcher, 1990). In Parkinson’s disease, SNr and GPi activity was posited to be increased through an imbalance in direct and indirect pathway striatal input, resulting in thalamic hypoactivity and consequent inability to initiate and perform movements (Albin et al., 1989; Alexander and Crutcher, 1990; DeLong, 1990).

More recently, the BGMT has become recognized more of an integration center in its own right and is considered to actively process synaptic input from multiple sources instead of just transmitting a rate code (Bosch-Bouju et al., 2013). In mice, a closed excitatory loop between VM and anterolateral premotor cortex (ALM) was found to be essential to allow movement initiation (Guo et al., 2017), and this loop could be gated by BG output (Catanese and Jaeger, 2020). The question of how in this integration framework the BGMT thalamus is engaged in mediating circuit dysfunction in Parkinson’s disease remains unanswered.

Thalamic neurons possess a strong T-type calcium current, which enables rebound bursting and could be involved in normal and pathological patterns of synaptic integration in rodents (Kim et al., 2017) and primates (Devergnas et al., 2016). Thalamic neurons are also heavily modulated through cholinergic and adrenergic input in their excitability state between waking and sleep modes (McCormick, 1989; McCormick and Prince, 1986). Given a near universal presence of homeostatic regulation of neural excitability in different neural circuits (Davis, 2006; Turrigiano, 2011), an increased inhibitory rate of BG input to BGMT during parkinsonian conditions is likely to engage such mechanisms, which could lead to an increase in excitability. In a series of elegant studies Bevan et al. showed that such homeostatic plasticity exists in the subthalamic nucleus in 6-OHDA lesioned mice where it counteracts reduced input from globus pallidus (Fan et al., 2012; Wilson and Bevan, 2011). This plasticity may be maladaptive in terms of motor function, however, and result in pathologically correlated activity (Chu et al., 2015; McIver et al., 2019). To test the hypothesis that BGMT neurons show changes in excitability in a parkinsonian condition, we obtained whole cell recordings from BGMT neurons in slices of adult mice and compared neuronal excitability between a control group and mice with unilateral 6-OHDA lesions. In support of our hypothesis we identified an increase in excitability of BGMT neurons in 6-OHDA lesioned mice, which was primarily due to a decrease in M-type potassium current. In the course of our studies we also for the first time characterized multiple aspects of intrinsic excitability of BGMT neurons in both normal and 6-OHDA lesioned mice that likely is essential in supporting closed loop thalamocortical excitation.

## Materials and Methods

### Animals

All animal procedures were approved by the Emory IACUC and adhered to the NIH Guide for the Care and Use of Laboratory Animals. For optogenetic stimulation and electrophysiological recordings, male and female C57BL/6J, Vgat-IRES-Cre mice (*Slc32a1*) aged 1-17 months (n=38) were used for terminal brain slice experiments. Of these mice 12 had been unilaterally 6-OHDA lesioned 1-10 months (mean period±SEM 4.7±0.8 months) before the experiment.

### Viral vector injections

Before surgery slow release buprenorphine SR (1 mg/kg) was administered subcutaneously to reduce pain as long-lasting analgesic. Mice were anesthetized with isoflurane (induction at 3–4% concentration and maintained at 1–2%) and head-fixed on a stereotaxic frame (Kopf Instruments). Ophthalmic ointment was applied to prevent corneal dehydration and a heating pad was used to maintain temperature at 37° C. A skin incision was made that allowed to perform craniotomies above ALM and SNr unilaterally on the right side of brain. To label GABAergic SNr terminals in BGMT with green fluorescence, 200-300nL of rAAV2/hsyn-EYFP was injected with a nano-injector (Nanoinject III, Drummond Scientific) at the average rate of 0.33 nL/s (total 300 nL during 15 mins) into the SNr targeting the Paxinos mouse atlas (Franklin and Paxinos, 2008) coordinates (in mm from Bregma): AP −3.2, ML 1.6, DV −4.4. To label ALM terminals in BGMT with red fluorescence and express ChR2 in the same terminals, rAAV2/CamkIIa-hChR2(T159C)-mCherry-WPRE was injected (at volume 300 nL) into the ALM cortex targeting the Paxinos coordinates (in mm from Bregma): AP 2.5, ML 1.5, DV −1.0 of some animals. This enabled us to subsequently selectively visualize the nigral and ALM terminal fields in BGMT and record from neurons within this field. After surgery, bacitracin ointment was applied to the region around the incision. Mice were weighed daily and assessed for health and comfort for 4 days post-surgery.

### 6-OHDA treatment

In a separate group C57BL/6J (n=11), and one Vgat-IRES-Cre mouse (*Slc32a1* were injected with 1 μL of 6-hydroxydopamine hydrochloride (6-OHDA) into the medial forebrain bundle (−1.2 AP, 1.2 ML, −4.75 DV) using the procedure described in Lundblad et al. (2004). Briefly, 0.02% ascorbic acid was added to a sterile saline solution (NaCl, 0.9% w/v) and 6-OHDA HCl powder (6-hydroxydopamine hydrochloride from Sigma or 6-OHDA hydrobromide from Tocris) was then dissolved to produce a final 6-OHDA concentration of 4.44 mg/ml. This method produced strong nigrostriatal lesioning as verified histologically through TH antibody staining (see Histology Methods Section).

### Slice preparation and solutions

On each experimental day, a mouse was deeply anesthetized with isoflurane and transcardial perfusion was performed with icy cold choline chloride solution containing (in mM): 117 ChCl, 2.5 KCl, 1.25 NaH_2_PO_4_, 26 NaHCO_3_, 10 dextrose, 0.5 CaCl_2_, 7 MgCl_2_, 1.0 sodium pyruvate, 1.3 L-ascorbic acid, which was bubbled with 95% O_2_-5% CO_2_. Mouse decapitation was performed with scissors, and the brain was quickly removed, immersed in cold choline chloride solution, and mounted on the flat surface of the microtome tray (Microm HM 650). Coronal thalamic slices (250 µm thick) were prepared and put to recover in a holding chamber in regular artificial cerebrospinal fluid (ACSF) at 32°C for 20 mins followed by room temperature. The ACSF contained (in mM): 124 NaCl, 2.5 KCl, 1.25 NaH_2_PO_4_, 26 NaHCO_3_, 10glucose, 2 CaCl_2_, 1.3 MgCl_2_. Chemicals were purchased from Sigma (St. Louis, MO, USA) or Abcam (Cambridge, MA, USA).

Individual slices were transferred to a recording chamber and continuously superfused with oxygenated ACSF at 26-28 °C at a flow rate of 2.5 ml/min. The BGMT region was visually identified by video microscopy (Olympus model BX51WI outfitted with differential interface contrast and an IR sensitive Dage MTI camera attached to the recording setup.) by mcherry expression due to cortical glutamatergic afferents projecting to BGMT neurons following AAV injection into ALM and by proximity to the mt fiber bundle. Neurons in slices without terminal label (n=1 mouse in 6-OHDA treated group, and n=17 mice in control group) were localized to BGMT according to matching their location with those find in labeled slices by proximity to the mt fiber bundle at the matching AP level (Franklin and Paxinos, 2008). Thereafter, whole-cell patch-clamp recordings were obtained at the soma under 60x magnification in BGMT neurons from control and 6-OHDA-lesioned mice with glass pipettes (at resistances 4-8 MΩ) pulled from 1.5 mm OD borosilicate glass on a Sutter P-97 puller (Sutter Instruments, Novato, CA).

### Determination of Intrinsic Excitability

To compare intrinsic excitability between 6-OHDA treated and control mice, we assessed I_h_, I_M_, I_T-Ca++_ current contributions to membrane potential trajectories. Current traces were recorded in whole-cell current clamp mode using a K-gluconate pipette solution containing (in mM): 130 K-gluconate, 10 NaCl, 10 KCl, 10 HEPES, 1 MgCl_2_, 0.5 Na-GTP and 1 Mg-ATP, 5 phosphocreatine, 0.1 spermine, 0.2 EGTA; titrated to pH 7.2 with KOH. The junction potential of this intracellular solution used for current-clamp with respect to the ACSF was calculated with JPcalc (Barry, 1994), and had a value of 14.2 mV. It was not subtracted from the measurements reported. Depolarizing and hyperpolarizing command current pulses at various duration and amplitude were injected into recorded somata via the patch pipette. To prevent spontaneous network firing, synaptic blockers of glutamatergic signaling DNQX (10 µM) and D-AP5 (50 µM) were included in the patch-clamp superfusion. A subset of neurons was exposed to the specific M-channel blocker XE-991dihydrochloride (10-20 µM) added to the bath after washing-in control ACSF to evaluate the role of M-type potassium channels in modulating 6-OHDA induced-hyperexcitability. While whole-cell configuration was established, each neuron was not stimulated for at least 5 minutes. Cells were included in the data if the resting membrane potential (V_*Rest*_) was at least −55 mV.

Data acquisition was performed using a Multi Clamp Amplifier 700B in conjunction with a customized LabVIEW (National Instruments) software interface. Whole cell patch clamp recordings were low-pass filtered at 10 kHz and digitized at 20 kHz. All analysis of electrophysiological data was performed using custom protocols using MATLAB (MathWorks, Inc.).

To evaluate intrinsic excitability, neurons were injected with depolarizing current pulses (ranging from 20 to 260 pA; 2000 ms in duration). The resting membrane potential of BGMT neurons typically was hyperpolarized enough (CON: −64.8 mV [-74.6,-60], n=19; 6-OHDA: −60.8 mV [-67.7,-49.0], n=17; Mann-Whitney test, p=0.0006; Figure 1E) so that a small depolarizing current injection would trigger a single low-threshold spike (LTS) burst. To deactivate the underlying T-type Ca^2+^ current, a bias current (CON: 59.8 pA [25.1, 246.7], n=19; 6-OHDA: 45.3 pA [-99.6, 150.5], n=17; Mann-Whitney test, p=0.04) was applied (CON: −57.2 mV [-64.1, −52.7], n=19; 6-OHDA: −54.8 mV [-69.2, −50.2], n=17; Mann-Whitney test, p=0.12) to achieve the desired membrane voltage (Jahnsen and Llinas, 1984a; Jahnsen and Llinás, 1984; Lundblad et al., 2004). The action potential firing frequency was calculated for each current step on top of the applied bias. Only action potentials that occurred at 50 ms or longer after the onset of the step current injection were included in the analysis (Dougherty et al., 2012). F-I curves (frequency of action potential firing as a function of injected current) were constructed. The rheobase was determined as the current amplitude at which a linear fit to the F-I curve evoked 3 Hz-action potential firing. To measure input resistance (R_*in*_), a hyperpolarizing −10 pA pulse current of 100 ms was applied and the voltage response amplitude was measured at 100 ms. The membrane time constant was determined based on voltage responses to −1nA current pulse injection of 0.5 ms duration, and calculated as elapsed time required for the evoked voltage response to decay back to 33% of the peak amplitude. The action potential voltage threshold (V_*T*_) was determined as the measured voltage where the value of dV/dt exceeded 10 mV/ms at the first AP in response to a 2000ms current step of minimal amplitude to elicit APs (Dougherty et al., 2012). To evaluate hyperpolarization-activated cationic currents (I_H_) hyperpolarizing current steps (range −200 to −50 pA; 50 pA increment, 2000 ms duration) were applied in current clamp mode at the resting membrane potential (mean±SEM; −62.7±0.8 mV; for CON & 6-OHDA all together). Percentage sag was measured as 100*(1 − V_ss_/V_peak_), where V_ss_ was the steady-state voltage deflection from baseline at 2000 ms after pulse onset, and V_peak_ was the peak negative voltage deflection from baseline (Narayanan and Johnston, 2007). To reveal effects of 6-OHDA treatment on rebound firing, hyperpolarizing current steps (range −500 to −50 pA, 50 pA increment, and 200, 500 or 2000 ms duration) were applied in current clamp mode. The baseline membrane potential at the time of applying steps on average was −56.5 mV [-63.7, −50] while a bias current [range: −17.5, 210.5 pA] was applied to stabilize a subthreshold voltage depolarized at this level to avoid T-type channel de-inactivation prior to pulse onset. For each individual voltage trace, the peak deflection and number of action potentials were calculated.

**Figure 1.**
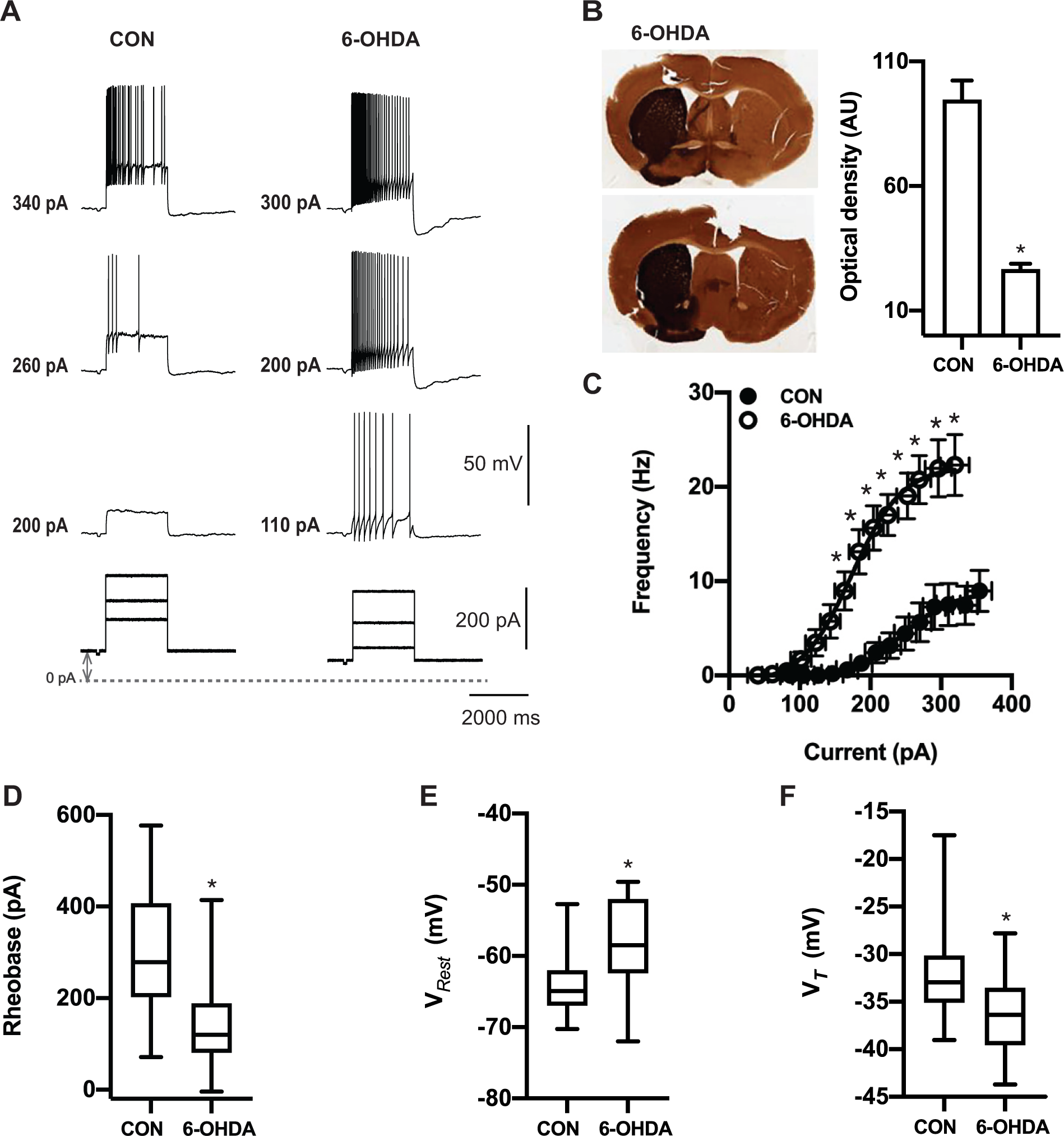
6-OHDA induced dopamine depletion elicits hyperexcitability in ventral motor thalamus region. **A**. Characteristic voltage responses to increasing current injections (bottom) recorded in a representative neurons of control mouse (CON, left upper panel) and of mouse 20 weeks after injection with 6-OHDA (right upper panel). To block glutamatergic and GABA-ergic synaptic inputs, DNQX (AMPA/kainite receptors antagonist, 10 µM), D-AP5 (NMDA receptor antagonists, 50 µM) and gabazine (10 µM) were applied in both situations. Dashed line designates the 0-pA level of bias current and up down grey arrow indicate bias current. In these examples baseline membrane potentials (CON: −65 mV and 6-OHDA: −67 mV) were depolarized up to −55 mV by applying bias currents (CON: 100 pA and 6-OHDA: 70 pA). **B**. Verification of successful 6-OHDA lesioning was achieved by tyrosine hydroxylase (TH) staining. *Left panel*. Representative digitized images of coronial striatal sections showing example tyrosine hydroxylase staining in control (left) and lesioned (right) hemispheres of two individual mice after administration 1 μl of 6-OHDA solution in the medial forebrain bundle. Top picture shows 68% decrease in TH immunoreaction of lesioned dorsal striatum by compared to the non-lesioned side (27.9 AU vs. 88.6 AU) four weeks after dopamine lost. At bottom, image of mice’s striatum lasting dopamine deficit for five months generating 70% reduction in TH staining (21.6 AU vs. 71.11 AU). *Right panel*. On average TH immunoreactivity calculated based on optical density was significantly higher at non-injected site compared to 6-OHDA treated site (CON vs. 6-OHDA side: 87.8±6.6 vs. 26.2±2.1, n=12; paired t-test, p<0.0001). **C**. Dopamine deficit for 5 - 40 weeks significantly enhanced firing frequency and shifts the F-I curve to the left (open circles) compared to controls (filled circles) (CON: n=19 vs. 6-OHDA: n=17; repeated-measures mixed-effects model, p≤0.015). The bias given to depolarize neurons up to −56.1 mV [-64.9, −50.5] to avoid T-type burst spiking was significantly higher for neurons from 6-OHDA treated than control mice (Mann-Whitney test, p=0.04; see method section). Each circle represents mean±SEM. **D**. For each individual neuron included in panel C, a Boltzmann sigmoidal fit was done based on injected current-firing rate relationship and tonic rheobase current that induced action potentials firing at 3 Hz was measured. Average tonic rheobase was significantly reduced by half after dopamine deficit (on right) compared to controls (CON: n=19 vs. 6-OHDA: n=17; Mann-Whitney test, p=0.0001). On average resting membrane potential was depolarized 6.4 mV higher in 6-OHDA treated (CON: −64.9 mV [-70.3, −52.7], n=46 vs. 6-OHDA: −58.5 mV [-72.0, −49.6], n=31; Mann-Whitney test, p<0.0001) (panel **E**). Action potential voltage threshold (V_*T*_) showed significant reduction compared to controls (CON: n=18 vs. 6-OHDA: n=17; Mann-Whitney test, p=0.014) (panel **F**, the same set of neurons as in panel **C** and **D**). Box and whisker plots represent medians, quartiles, and 5-95% percentiles.

### Statistical Analysis

Statistical analysis was performed using Prism 6 (GraphPad Software, La Jolla, Ca, USA). Statistical significance between comparisons of two groups was achieved using paired or un-paired t-test for data that was normally distributed (as measured by either Pearson or Shapiro-Wilk normality test). When data was not normally distributed, Wilcoxon matched-pairs signed rank tests or unpaired Mann-Whitney U tests were administered. Comparisons of F-I curves between control and 6-OHDA treated mice were performed using two-way repeated-measures ANOVA on spike rate by injection level, and mixed-effect repeated measures. Normally distributed data is presented as mean±SEM in the text; whereas skewed data is presented as medians with quartiles. Statistically significant differences are represented on figures by asterisks.

### Immunohistochemistry for tyrosine hydroxylase and quantification of nigrostriatal innervation

Coronal sections (150-200 μm, obtained during preparation of slices for electrophysiology) were submerged in 4% paraformaldehyde in 0.1M PB. The sections were then embedded in 0.5% gelatin in distilled water, and the gelatin block was immersed again in 4% paraformaldehyde overnight. The tissue was then sectioned to 40 mm coronal sections using a vibrating microtome (Leica) the gelatin embedded sections were cut into 40 mm sections and stored at −20°C in antifreeze solution. To verify denervation of the nigrostriatal pathway induced by 6-OHDA treatment, sections at the level of the striatum were pretreated with 1% normal goat serum, 1% bovine serum albumin and 0.3% Triton X-100, and then incubated in rabbit anti-tyrosine hydroxylase (TH) antibody solution (Millipore cat # AB152, 1;300) overnight. This was followed by incubation in secondary biotinylated antibodies, then in Avidin-biotin-peroxidase complex (ABC) solution (1:200; Vectastain standard kit, Vector) for 90 min. The sections were then placed in 0.025% 3-3’-diaminobenzidine tetrahydrochloride (DAB, Sigma-Aldrich, St. Louis, MO), 0.01M Imidazole (Fisher Scientific, Pittsburgh, PA) and 0.006% H2O2 for 10 min. All incubations were done at room temperature. The sections were mounted on slides, cover-slipped, and digitized with an Aperio Scanscope CS system (Leica).

To quantify the loss of nigrostriatal innervation, the optical density of TH-stained areas was measured in the striatum of the lesioned hemisphere of 6-OHDA treated animals, and these values compared against the striatal optical density from the contralateral hemisphere of the same slice. ImageJ. The scanned images were corrected for brightness, converted into 16-bit grayscale format and inverted. For each animal, measurements of the optical density were obtained in the dorsolateral striatum in two sections (approximately at anteroposterior planes −0.7 mm and 0.1 mm from Bregma, according to Paxinos and Franklin, 2001). To control for differences in background staining, the optical density measured in the corpus callosum was subtracted from the striatal measurements.

### Computer Simulations

The simulations were carried out with an integrated NEURON (v.7.7) + Python (v.3.7) environment (Hines and Carnevale, 1997), with an adaptive time-step integration, i.e., Cvode solver. The simulated temperature was 27° C as the average value given by the experiments. The model source code will be publicly available on the Senselab ModelDB (http://senselab.med.yale.edu) and GitHub (https://github.com/FrancescoCavarretta/VMThalamocorticalNeuronModel) upon publication.

The Thalamocortical Cell (TC) was implemented as a multicompartmental biophysically-detailed model that replicates the full dendritic tree and firing behaviors that characterize the TCs of the BGMT thalamus.

### Morphology

We used a morphological reconstruction of a ventromedial (VM) nucleus TC belonging to the Janelia MouseLight Dataset (https://www.janelia.org/project-team/mouselight; id. AA0136). Although these VM morphologies included full reconstructions of the axon, we retained only the initial 70 µm of it, which is putatively the axon initial segment (AIS). Therefore, the morphology of our TC model was comprised of three subcellular sections, the soma, the dendrites, and the AIS. The experimental procedure used for reconstructing the MouseLight morphologies did not allow to measure the diameters of the neurites. We therefore fixed the diameter of the AIS to 1.5 µm; the soma was replaced with a single compartment of 293 µm^2^ in surface (Sawyer et al., 1989); and we defined a model for the dendritic diameters based on the 2/3 Rall’ Power Law, of which parameters were directly estimated from 8 reconstructions of BGMT TCs from the physiological data set presented in this study.

### Membrane Properties

The passive parameter values were given by a uniform specific membrane resistivity (rm) of 26.0 MΩcm2, cytoplasmic resistivity (ri; i.e., intracellular or axial resistivity) of 60.0 Ωcm, specific membrane capacitance (cm) of 1.0 µF/cm^2^ (Gentet et al., 2000), and resting potential (Vrest) of −75.75 mV. To simulate the firing behavior of BGMT TCs, we distributed 10 classes of active membrane conductances along the TC morphology (Table 1). The active conductances were NaF (i.e., sodium channels that are not Nav1.6 types), NaP (i.e., Nav1.6 type), KDR, KA, I_H_, CaT, CaL, SK (i.e., small-conductance calcium-activated sodium channels), I_M_ (i.e. Kv7), and ANO2-CaC channels (i.e., calcium-activated chloride channels). The models of NaF, KDR, KA, I_H_, CaT, CaL, and SK channels were imported from a previously published model of ventrobasal thalamocortical neurons (Iavarone et al., 2019). To match our measures of the sag amplitude, we shifted the inactivation curve by −11 mV for the I_H_ channel.

**Table 1.** Subcellular distributions of ion channels in thalamocortical neuron model. Number given in units of S/cm^2^

| Channel | AIS |  | Soma | Dendrites |  |
| --- | --- | --- | --- | --- | --- |
|  | Distal | Proximal |  | 1<sup>st</sup> order | 2<sup>nd</sup> order or above |
| NaF | 0.008 | 0.152 | 0.008 | 0.008 | 0.004 |
| NaP | 0.0019 | 0.0001 | 0.0001 | 0.0001 | 0.0001 |
| K<sub>DR</sub> | 0.08 | 1.52 | 0.08 | 0.08 | 0.08 |
| KA | 0.005 | 0.095 | 0.005 | 0.005 | 0.005 |
| KM | 0.0165 | 0.00033 | 0.00033 | 0.00033 | 0.00033 |
| CaL | 0.0041 | 0.0041 | 0.0041 | 0.00205 | 0.00205 |
| SK | 0.0008 | 0.0008 | 0.0008 | 0.0004 | 0.0004 |
| ANO2-CAC | 0.00275 | 0.00275 | 0.00275 | 0.001375 | 0.001375 |
| CaT | 0 | 0 | 0.000085 | 0.00017 | 0.0000425 |
| I<sub>H</sub> | 0 | 0 | 0.000035 | 0.000035 | 0.000035 |

This alteration made the sag amplitudes increasing monotonically with hyperpolarizing currents between −50 pA and −200 pA. Indeed, without this alteration, the sag amplitude was lower with current injection of −200 pA than −150 pA. We derived the NaP from the NaF shifting the activation and inactivation curves by 14 mV (Hu et al., 2009). We based the I_M_ channel on experimental data of neocortical pyramidal cells (Battefeld et al., 2014), adding a constant term (of 0.025) to the equation of activation (m):

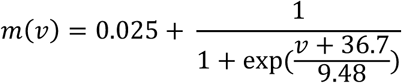

This alteration of the original equation increased the opening fraction of I_M_ channels, which decreased the Vrest of the TC model to our physiological values. We implemented a model of ANO2-CaCC channel based on the experimental data of [Pifferi et al., 2009], using a reversal potential of −86 mV. This channel is responsible for early firing rate adaptation in TC neurons (Ha et al., 2016). The ionic subcellular distribution of each ion channel resulting in a good match with the physiological recordings based on manual tuning are given in Table 1. In the TC model, intracellular calcium was structured in three separate microdomains with distinct decay constants. In particular, calcium flowing through CaL channel bound with two microdomains with different percentages (25.8% and 74.2%), activating SK and ANO2-CACC channels (decay time constant: 14 ms and 50 ms, respectively). Instead, calcium flowing through CaT channel bound with a third microdomain (decay time constant: 50 ms), without activating ion channels (Womack et al., 2004).

### Analysis of model cell properties

To compare the passive cell properties of our BGMT TC model to our physiological recordings, we simulated somatic current injections. Simulations were necessary for their measurements, as they resulted of the non-linear combination between morphological features, passive properties, along with the configuration and the dynamics of the active membrane conductances. To measure the input resistance (Rin), we simulated a hyperpolarizing injection of −10 pA for 100 ms, calculating the Rin as the deviation from the resting potential, measured at 100 ms, divided by the current intensity. We thus estimated an Rin of 218.2 MΩ. To measure the membrane time constant (τ), we used a double exponential fitting (a·exp (t/τ1)+ c ·exp (t/τ2)) to the decaying phase of the membrane voltage observed with a hyperpolarizing current pulse of −1 nA (0.5 ms). The membrane time constant then corresponded to the time constant of the slowest exponential term. We thus estimated that τ was 19.1 ms for our TC model, consistent with our experimental measures. The cell capacitance (C) was calculated as the ratio between τ and R_in_, and resulting in an estimate of 87.4 pF, consistent with our experimental measures. Finally, the Vrest was −77.3 mV, which was consistent with our experimental measures of −65 mV given the junction potential of −14.2 mV.

## Results

The goal of our study was to determine any changes in neural properties in the area of motor thalamus that receives BG input (BGMT) in mice unilaterally treated with 6-OHDA in the median forebrain bundle as a standard rodent model of robust dopamine neuron lesioning. To this end, we obtained brain slice recordings of BGMT neurons from a total of 26 control mice and of 12 6-OHDA treated mice. The BGMT was either defined through fluorescent label of nigral GABAergic input or through location with respect to the mt fiber bundle. We broadly ascertained excitability properties that govern intrinsic neural dynamics in these cells that could be altered in Parkinsonian states due to homeostatic plasticity mechanisms.

### Changes in firing frequency – current (f/I) relationships

Thalamic neurons are well known to have two distinct firing modes, often called burst and tonic firing (Ramcharan et al., 2000). Burst firing is enabled when the membrane is sufficiently hyperpolarized to de-inactivate a T-type calcium current (CaT) (Gutierrez et al., 2001; Jahnsen and Llinas, 1984b; Llinas and Steriade, 2006). In awake animals, tonic firing generally predominates due to the depolarizing baseline of synaptic inputs and cholinergic modulation (McCormick and Prince, 1986; Sherman and Guillery, 2002). To compare the tonic firing properties of BGMT neurons between control and 6-OHDA treated conditions we therefore depolarized whole cell recordings to a level of −55 mV with a tonic bias current to inactivate CaT, and added positive current injection pulses on top of this bias current to ascertain the minimum current needed to elicit tonic firing (rheobase), the spike threshold (V_T_), and the relationship between injected current amplitude and firing frequency (F-I) (Fig. 1). We found a significant difference in all these parameters for neurons obtained from 6-OHDA treated animals (n=17 neurons) compared to neurons obtained from controls (n = 19 neurons). BGMT neurons from 6-OHDA treated mice showed a significantly lower rheobase (Fig. 1D) and a significantly lower spike voltage threshold (Fig. 1F), and finally a dramatically higher spike frequency with increasing current injection amplitudes (Fig. 1C) than neurons from control mice. Without bias current injection, neurons from 6-OHDA treated mice additionally showed a 6.5 mV more depolarized resting membrane potential than neurons from control mice. These results show our main finding that BGMT neurons in 6-OHDA treated mice are considerably more excitable by depolarizing input than neurons in control mice.

### Passive Properties of BGMT neurons in control and 6-OHDA lesioned mice are similar

We next asked the question of whether any changes in passive properties of BGMT neurons in 6-OHDA lesioned mice might account for the increase in excitability. For example, if such neurons were to shrink in size, they would show a lower capacitance and an increased input resistance to make them more excitable with a given amount of current injection. We found, however, that passive properties remained unchanged. Measures of the membrane time constant (Fig. 2A), membrane capacitance (Fig 2B), and input resistance (Fig 2C) (see Methods for details) revealed no significant differences between control and 6-OHDA lesioned mice. Membrane capacitance and input resistance were distributed between a similar range of values for both conditions and showed an inverse relationship as expected (Fig. 2D). We also plotted the respective distributions of membrane time constant, (Fig. 2E) membrane capacitance (Fig. 2F) and input resistance (Fig. 2G).

**Figure 2.**
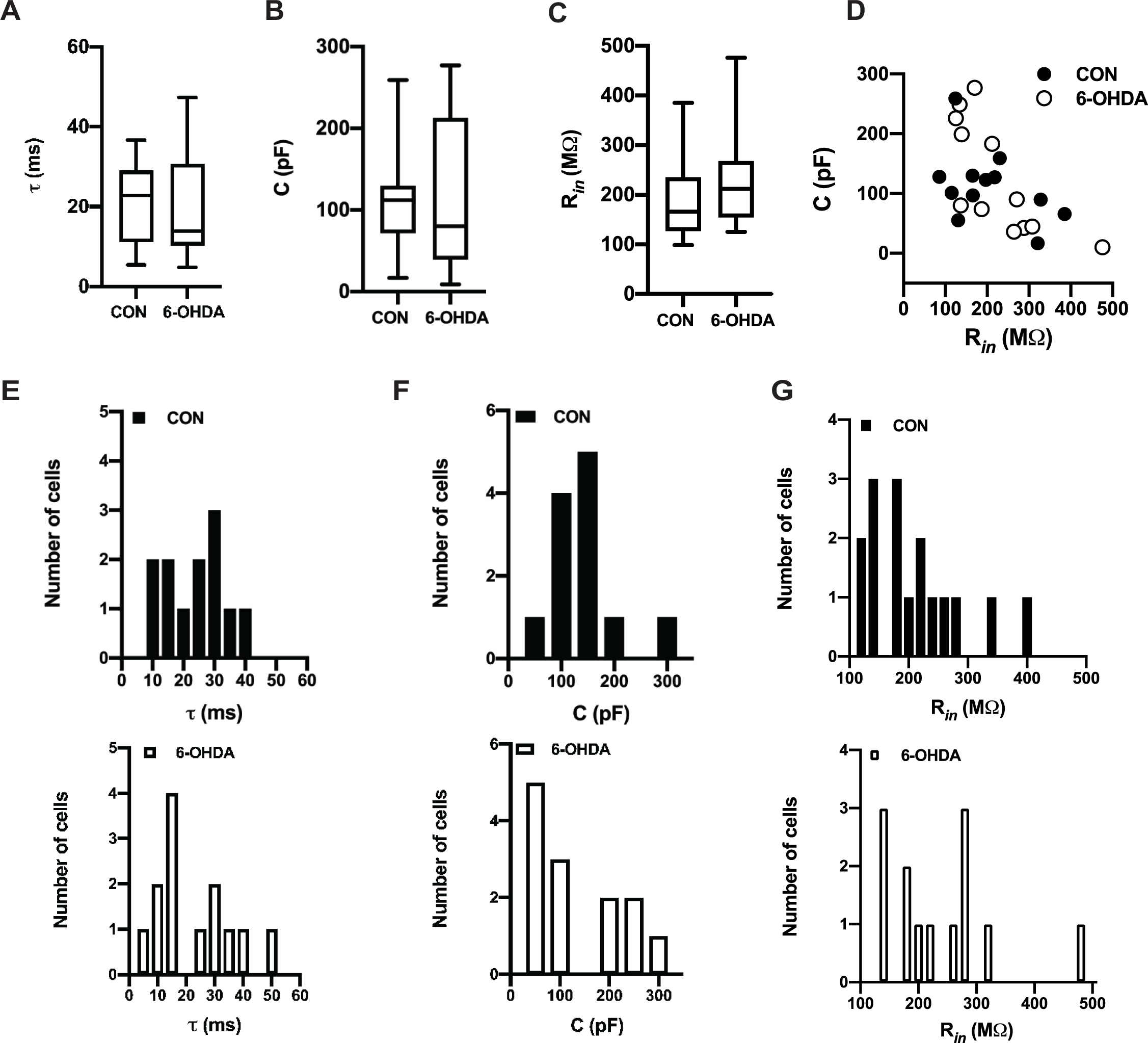
Resting membrane properties remaining intact after hydroxy-dopamine injection. Electrophysiological passive properties for the data presented in Figure 1. A-C. Dopamine depleted hyperexcitable neurons did not shown significant changes in time constant τ (CON: 22.8 ms [5.4, 36.6], n=12; 6-OHDA: 13.8 ms [4.8, 47.2], n=13; Mann-Whitney test, p>0.99), membrane capacitance (CON: 112.0 pF [16.8, 259.1], n=12; 6-OHDA: 80.2 pF [8.9, 277.0], n=13; Mann-Whitney test, p=0.65) and input resistance measured at resting membrane potential (R_in_: CON: 166 MΩ [98.5, 385.0], n=17; 6-OHDA: 212.0 MΩ [125, 476], n=13; Mann-Whitney test, p=0.19). **D**. In 6-OHDA mouse model, relationship between membrane capacitance and R_in_ is comparable to healthy cells in motor thalamus area. Because capacitance linear relationship with cell surface area pointing no changes occur in surface area. Box and whisker plots represent medians, quartiles, and 5-95% percentiles. **E-G**. Histograms of data presented in panel A-C for controls (on top) and after treatment (bottom). Individual histograms represent distribution of τ, capacitance and input resistance data based on bin width 5ms, 50 pF and 20 MΩ respectively.

### 6-OHDA treated mice show an increase in sag with hyperpolarizing current injection in a subpopulation of neurons

The hyperpolarization activated (I_H_) current associated with cyclic nucleotide gated (HCN) channels has been shown to counteract inhibitory input in subthalamic neurons (Atherton et al., 2010), but also can limit excitatory input (Sheets et al., 2011) and stabilize the membrane potential to prevent bistability (Williams et al., 2002). Further, alteration in I_H_ has been implicated in several disease models, including Alzheimers (Eslamizade et al., 2015), and epilepsy (Noam et al., 2011). A hallmark of I_H_ is that it induces a “sag” in the response of the membrane potential to hyperpolarizing current steps. We determined sag responses in BGMT neurons (see Methods), and found that most but not all (Fig. 3D) recorded neurons showed a sag that was blocked by the selective I_H_ blocker ZD7288 (Fig. 3C). Comparing recordings from control and 6-OHDA lesioned mice we found that the sag amplitude of neurons that did show a sag response was larger in BGMT neurons from 6-OHDA treated mice than in controls (Fig. 3A,B). The increase was moderate, reaching significance for injection amplitudes of −50 and −100 pA, but barely missing it for −150 and −200 pA current injection steps (Fig. 3B). Since BGMT neurons show T-type calcium current dependent rebound bursts (Edgerton and Jaeger, 2014) similar to subthalamic neurons, the role of such an increase may be analogous to that found by Atherton et al (2010), and decrease the propensity for burst firing with strong GABAergic BG input transients.

**Figure 3.**
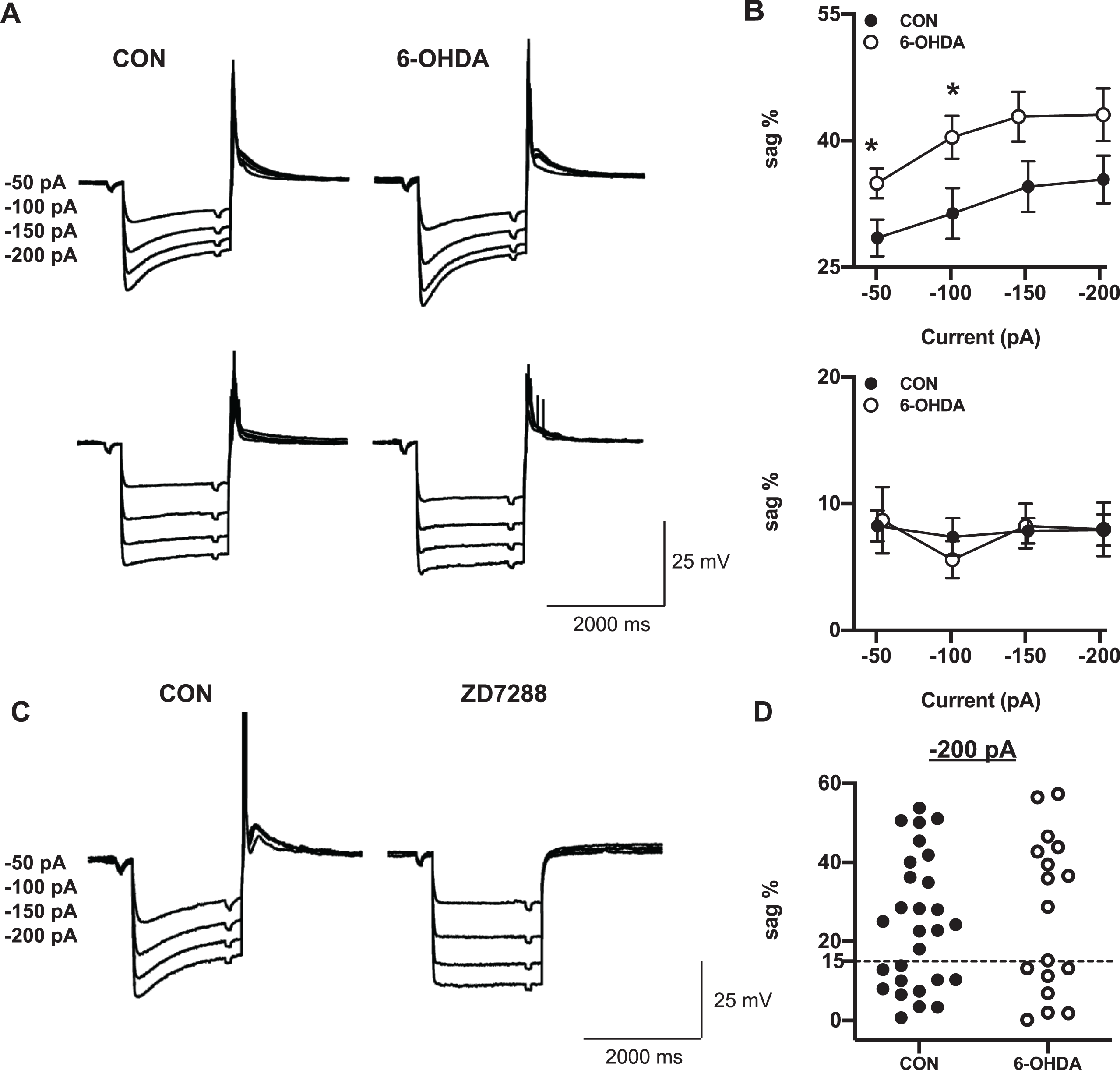
Sag amplitude is increased after 6-OHDA treatment. **A.*Top***. Responses to hyperpolarizing current steps (ranging from −200 to −50 pA, 50 pA increment) show the typical “sag” resulting from activation of I_H_ in a majority of neurons. Some neurons, however, showed little or no sag. We defined a 15% sag amplitude (difference between peak hyperpolarization and subsequent steady state potential – see Methods) as a threshold for neurons showing a discernible sag. The panels depict grand averages of voltage responses from cells with > 15% sag responses in control mice (n=16 neurons; Vrest = −61.0±0.78 mV) and 6-OHDA lesioned mice (n=8 neurons; Vrest = −60.9±1.01 mV). The average sag was more prominent 2-10 months following 6-OHDA injection (right panel) than in controls (left panel). ***Bottom***. Panels depicting the grand averages of responses to the same current injections in neurons with no discernible sag (control: n=9 neurons; Vrest = −67.0±1.39 mV; 6-OHDA treated: n=6 neurons; Vrest = −61.6±3.5 mV). **B.*Top***. The plots show averages of sag% as a function of injected step size for the neurons with >15% sag shown in panel A (top). 6-OHDA treatment resulted in an about 10% increase in “sag” magnitude compared to controls, which was significant for −50 and −100 pA current steps, and narrowly missed significance for −150 and - 150 pA current steps (CON vs. 6-OHDA: −50 pA : 28.5±2.2% (n=16) vs. 35.0±1.8% (n=8), paired t-test, p=0.03; −100 pA : 31.4±3.0% (n=17) vs. 40.4±2.5% (n=9), p=0.03; −150 pA : 34.6±3.0% (n=16) vs. 42.8±2.9% (n = 8), p=0.06; −200 pA : 35.4±2.8 (n= 17) vs. 43.1±3.1 (n = 9), p=0.08). Each circle represents mean±SEM. ***Bottom***. Same plots for neurons with <15% sag amplitude (average traces shown in panel A bottom). In these neurons, the amplitude of the remaining sag was not dependent on injection step size, and was not different between controls and 6-OHDA treated mice (CON vs. 6-OHDA: −50 pA : 8.2±1.2% (n=10) vs. 8.7±2.6% (n=7); −100 pA : 7.4±1.5% (n=11) vs. 5.6±1.5 (n=7); −150 pA : 7.8±1.0 (n=10) vs. 8.2±1.7(n=7); −200 pA : 7.9±1.2 (n=11) vs. 7.9±2.1 (n=8); mixed effect test, p≥0.4). **C**.Characteristic membrane potential responses to hyperpolarizing current steps in a representative control neuron. To block I_H_ currents, external ZD7288 (specific I_H_ antagonist) was added; which resulted in a complete elimination of the sag response, indicating that it was indeed due to I_H_. **D**.Distribution of sag amplitudes in voltage responses to 2 s −200 pA currents steps for different neurons. The dashed horizontal line designates the division into neuronal populations with no discernible sag - (sag<15%) and clear sag expressing cells (sag>15%). Note that the distribution especially for the 6-OHDA treated population is clearly bimodal.

### T-type Ca channel dependent rebound burst firing is increased in neurons from 6-OHDA treated mice

We next directly addressed the question whether T-type calcium current mediated rebound bursts were affected in BGMT neurons in 6-OHDA lesioned mice. To elicit rebound bursts, we injected negative current steps for 0.2, 0.5, or 2 s with amplitudes of between −50, and −500 pA (Fig. 4). As expected from T-type current mediated rebound bursts in thalamic neurons and shown in our previous study (Edgerton and Jaeger, 2014) these hyperpolarizing steps elicited strong 2-8 action potential bursts riding on a broader peak of depolarization caused by I_T_ (Llinás and Jahnsen, 1982). As expected, longer and stronger hyperpolarizing steps elicited stronger rebound bursts as measured by the number of action potentials in the rebound (Fig. 4E-F). In contrast to an attenuation of rebounds that might be expected from an increase in I_H_ (Atherton et al., 2010), however, we found a significant increase in the number of rebound burst spikes in BGMT neurons from 6-OHDA lesioned mice for all stimulus conditions except the smallest and shortest step depolarization (Figure 4D-F). A typical example for a BGMT neuron from a control and a 6-OHDA lesioned mouse is shown in Fig. 4A-C.

**Figure 4.**
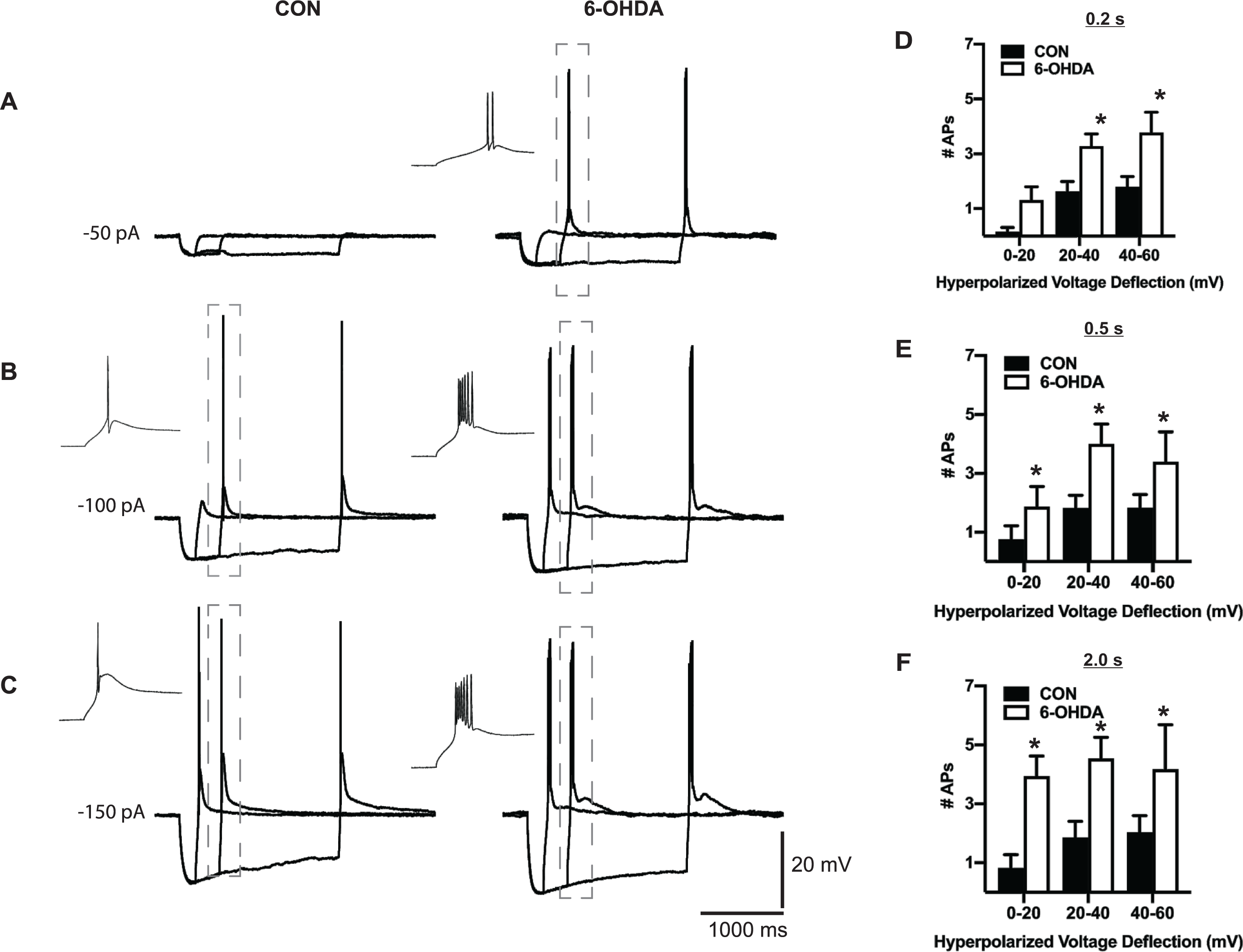
BGMT neurons from 6-OHDA treated mice showed stronger rebound burst firing. **A-C**. Single BGMT neuron voltage traces from representative control (left) and 6-OHDA (right) mice showing responses to 50, 100, 150 pA hyperpolarizing stimulus steps at durations of 0.2, 0.5 or 2 s. Dashed rectangles mark rebound bursts expanded in the inset windows to the left. **D-F**. Bar graph pairs represent the average number of action potentials per rebound burst ± SEM in controls (black bars) and in 6-OHDA treated mice (white bars). Data were sorted by the amplitude of hyperpolarization reached at the end of the step current hyperpolarizing current pulse (step amplitudes ranged from −50 to −500 pA at 50pA increments). Using this measure instead of the current injection amplitude avoids conflating rebound properties due to voltage-dependent channel de-inactivation with different levels of hyperpolarization reached due to passive input resistance of differently sized neurons. Prior to step current injection, a tonic bias current (CON: 90.7±15.7 pA; 6-OHDA: 54.1±16.07 pA) was adjusted to maintain the membrane potential at a similar level for all neurons (average of −56.52±0.6 mV), again in order to test for rebound properties under comparable voltage levels. The results show that the number of APs was increased up to 80% in neurons from 6-OHDA treated mice, which was significant for almost all stimulus conditions (0.2 s step: CON n= 15 vs. 6-OHDA n= 17; mixed-effects test, p=0.2, p=0.009, p=0.0008; 0.5 s: CON n= 12 vs. 6-OHDA n= 12; mixed-effects, p=0.05, p=0.004, p=0.03; 2.0 s: CON n= 12 vs. 6-OHDA n= 14; mixed-effects, p=0.005, p=0.001, p=0.002).

### The increase in tonic spiking frequency after 6-OHDA treatment is at least partly explained via a reduction of I_M_ K^+^ channels in BGMT neurons

A previous study in ventrobasal (VB) corticothalamic neurons showed that the muscarine sensitive potassium current (I_M_) carried by Kv7.2 and Kv7.3 channels can lead to a decrease in Vrest, a decrease in firing frequency with depolarizing current steps, and a decrease in burst spikes at the offset of hyperpolarizing steps (Cerina et al., 2015). Since these observations fit the profile of our observed changes after 6-OHDA treatment quite well, we wondered whether they are associated with a decrease in I_M_ current. We used the specific KV7 blocker XE-991 (Greene et al., 2017) to determine whether blocking KV7 current has a differential effect on BGMT neurons from control or 6-OHDA treated mice. We found indeed that application of 10-20 mM XE-991 resulted in a marked increase in tonic firing frequency in response to depolarizing steps in neurons from control mice (n=8 neurons, Fig. 5A,C), indicating the presence of KV7 channels and a role in dampening tonic spike frequencies. In addition, XE-991 in neurons from control mice showed a significant lower rheobase (Fig. 5D), and lower spike threshold (Fig. 5E). In contrast, none of these effects were observed in neurons from 6-OHDA treated mice (n=10 neurons), The lack of effect of XE-991 indicates a lack of functional I_M_ current in these neurons. Together, these results indicate that a reduction in I_M_ current can likely account for the majority of changes seen in intrinsic excitability in BGMT neurons following 6-OHDA treatment.

**Figure 5.**
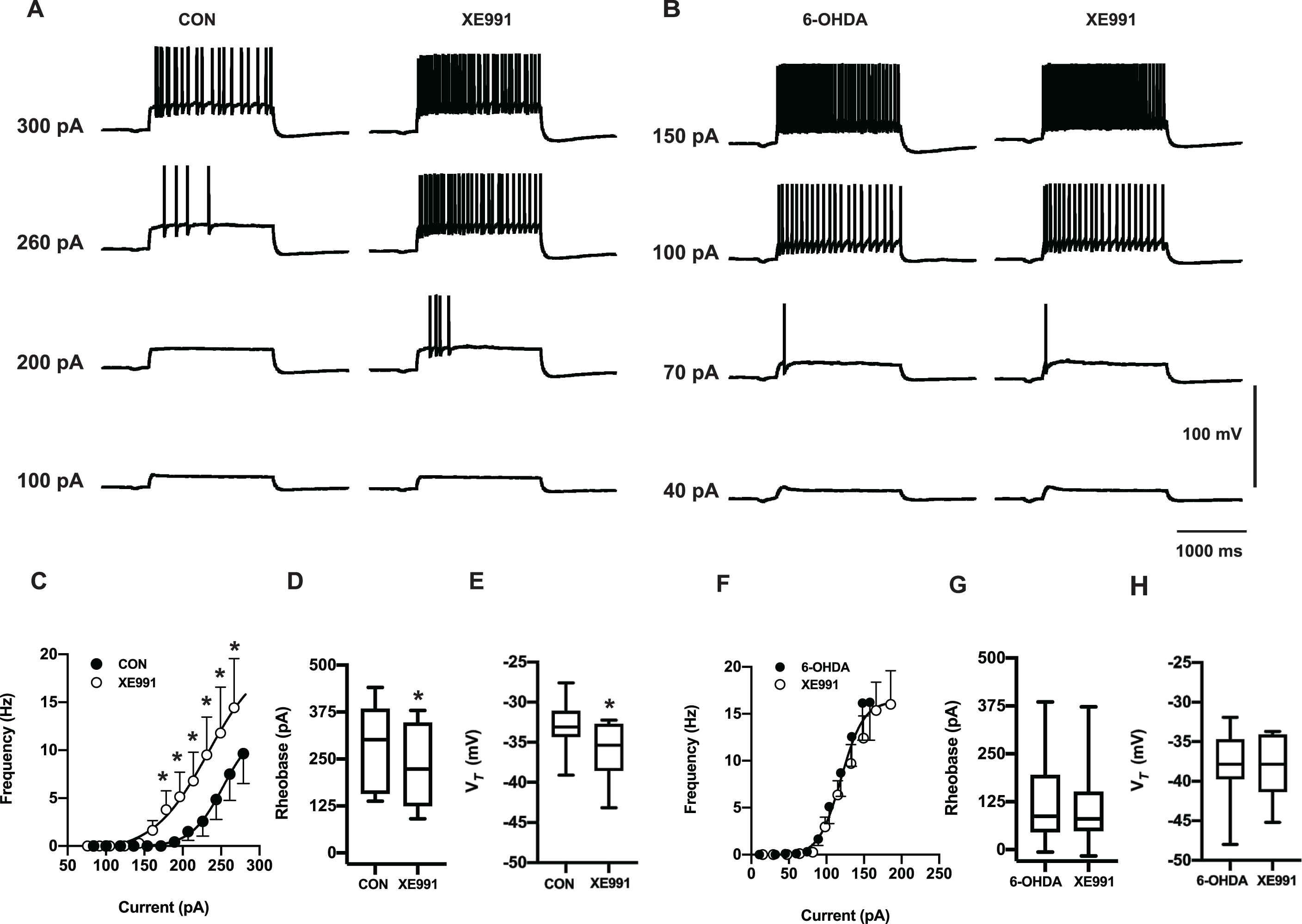
6-OHDA application enhances tonic spike firing in BGMT neurones via a reduction of I_M_ K^+^ conductance. **A**. Voltage responses to increasing depolarizing step current injections (100, 200, 260, 300 pA) for a representative neuron of control mouse before (left panel) and after 10 minutes of adding 10-20 µM XE-991 to the bath ACSF(right panel). To block ionotropic glutamatergic and GABA-ergic synaptic inputs, DNQX (AMPA/kainate receptors antagonist, 10 µM), D-AP5 (NMDA receptor antagonists, 50 µM) and gabazine (GABA-A antagonist, 10 µM,) were present in the ACSF throughout. **B**. Voltage responses to increasing pulse current injections (at amplitude 40, 70, 100, 150 pA) of respresenative neuron from 6-OHDA treated mouse before (left panel) and after (right panel) 10-mins application of XE-991. **C**. 10-20 mins bath exposure to XE-991 (10-20 µM) enhanced firing frequency in untreated control neurons and shifted the F-I curve to the left (CON vs. XE-991: n=7; two-way repeated-measures ANOVA, p≤0.02), suggecting I_M_ inhibition induced hyperexcitability. Baseline membrane potentials were depolarized up to −58.4±1.2 mV by applying bias current 83.9±26.1 pA in order to inactivate T-type Ca^2+^ currents and prevent burst firing. Each circle represents mean±SEM. **D**. For each individual neuron included in panel C, a Boltzmann sigmoidal fit was done for the I-F curve and the intercept at 3 Hz firing was determined as rheobase (see Methods). Neurons after exposure to XE-991 shown significant reduction in the rheobase compared to control ACSF (CON: 301.5 pA [137.7, 440.7]; CON + XE-991: 223.3 pA [90.6, 379.2], n=7; Wilcoxon test, p=0.015). **E**. Average voltage threshold (V_*T*_) significantly decreased in presence of XE-991 (CON: −33.1 mV [-39.1, −27.6]; CON + XE-991: −35.4 mV [-43.1, −32.2]; Paired t-test, p=0.005). **F-H**. XE-991 did not generate further changes in firing frequency of neurons from 6-OHDA treated mice (6-OHDA vs 6-OHDA + XE-991, n =10; repeated-measures mixed-effects model, p>0.24) in mice unilateraly lesioned 1-7.1 months (mean period±SEM: 4.0±1.0 months). Neither tonic rheobase (6-OHDA: 87.1 pA [-6.5, 385.5]; 6-OHDA + XE-991: 80.0 pA [-16.6, 372.7], n=10; Wilcoxon test, p=0.08) and V_*T*_ (6-OHDA: −37.8 mV [-48,-31]; 6-OHDA + XE-991: −37.8 mV [-45.2,-33.7], n=10; Paired t-test, p=0.88) showed changes in the presence of XE-991. Box and whisker plots represent medians, quartiles, and 5-95% percentiles.

### Analysis of I_M_ current effects in a biophysically realistic neuron model

We simulated the current step protocols as used in our experimental approach. Reducing or blocking the I_M_ current resulted in increased tonic firing as well as a lower rheobase (Fig. 6A,B). Full blockade of the I_M_ conductance also resulted in an increase in rebound burst spikes following hyperpolarizing current steps, while a reduction to 50% was not sufficient to cause this effect (Fig. 6C,D). Interestingly, a full blockade of I_M_ could also increase the sag amplitude when we simulated the hyperpolarizing steps used in our study (Fig. 6 E,F) however, this effect by itself was not large enough to explain the magnitude of differences seen between control and 6-OHDA lesioned mice in our slice recordings. Therefore, an additional increase in I_H_ current amplitude is likely to be required in order to match our experimental findings (Fig. 6F, No I_M_, 300% I_H_).

**Figure 6.**
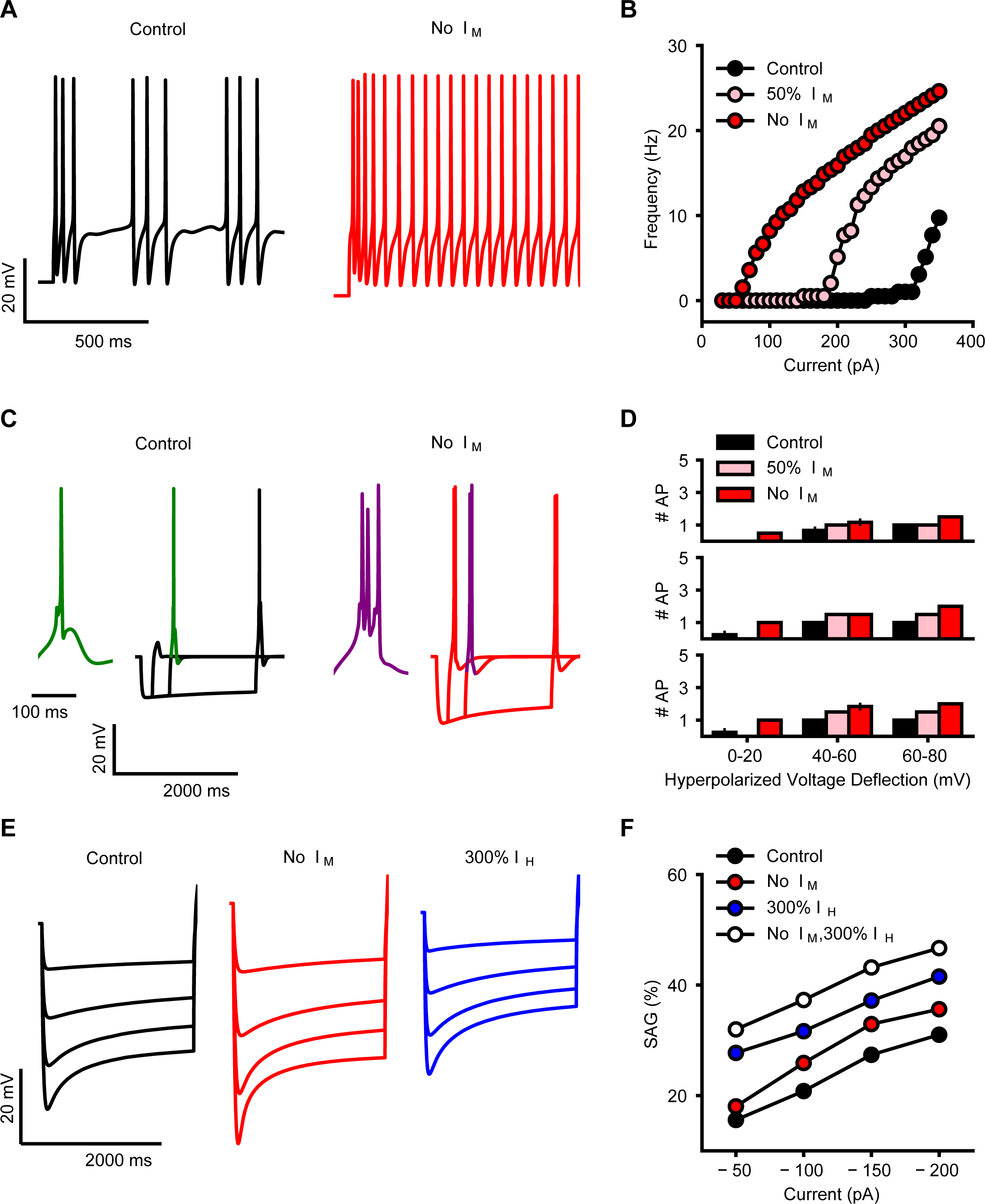
**A**. Simulated firing of BGMT thalamocortical neuron to somatic current injection (350 pA; 2s of duration) for control (black trace) and after I_M_ blockade (red trace). For both, membrane potential was held at ∼-64 mV to inactivate T calcium channels and evoke tonic firing upon depolarization. I_M_ blockade increases the model firing rate (compare black and red traces). **B**. Injected current-firing rate relationship of the model (F-I plot) for control (black), after 50%-reduction (pink) and full blockade (red) of I_M_ current, with current intensities of 30-360 pA. I_M_ reduction and blockade shifts the F-I curve leftward, resulting in a decrease in spike threshold as well as an increase in firing rate. **C**. Rebound bursts in the model neuron evoked by hyperpolarizing current steps (−100 pA) of different durations (200 ms, 500 ms, 2000 ms), for control (black) and after I_M_ blockade (red). Insets show an expanded waveforms of a burst (green, purple). Membrane potential was held at ∼-71 mV. I_M_ reduction increases the rebound burst duration, i.e., spike count (compare green and purple wave form). **D**. Spike count in rebound bursting as in C, evoked by current steps of 200 ms (top), 500 ms (center), and 2000 ms (bottom), for control (black), after 50%-reduction (pink) and full blockade (red) of I_M_ conductance. **E**. Sag generated in response to hyperpolarizing current pulses of increasing intensities (50-200 pA; no holding current) for control (black), after I_M_ blockade (red), and 200%-increase of I_H_ (blue). **F**. Sag amplitude vs hyperpolarizing current intensity for control (black), after I_M_ blockade, 200%-increase of I_H_ (blue), or combination of both changes (white). I_M_ blockade makes a small increase in sag amplitudes (compare red and black), while I_H_ increase brings the amplitudes to similar levels as observed in 6OHDA-lesioned condition (compare blue, white and black), combination of both changes allows to replicate the increase in the sag observed experimentally (compare white and black).

## Discussion

Changes in cellular properties have been observed in multiple key structures related to Parkinsonian motor dysfunction beyond primary striatal dysfunction directly elicited by dopamine depletion. Notably, in an elegant set of studies the Bevan lab showed that in the subthalamic nucleus (STN) 6-OHDA treatment in rodents leads to changes in synaptic as well as intrinsic properties (Chu et al., 2017; Chu et al., 2015; Fan et al., 2012; Kovaleski et al., 2020; McIver et al., 2019; Wilson and Bevan, 2011). A prominent synaptic change in STN is given by homeostatic upregulation of the GPe GABAergic input conductance through a mechanism caused by cortical NMDA input overactivation (Chu et al., 2015), which contributes to the emergence of pathologically correlated STN – GPe activity patterns (Magill et al., 2001; Mallet et al., 2008; Walters et al., 2007). In addition, NMDA overactivation after 6-OHDA depletion results in a downregulation of STN autonomous spiking due to an increase in K_ATP_ channel activity (McIver et al., 2019). These studies demonstrate that pathological input levels and input patterns can lead to homeostatic changes in cellular properties without the need for direct dopamine modulation. The BGMT in rodents is very sparsely if at all innervated by dopaminergic fibers (García-Cabezas et al., 2009), making direct dopaminergic effects unlikely. However, BG output from SNr has been shown to be more synchronized and bursty following 6-OHDA treatment in rodents (Avila et al., 2010; Brazhnik et al., 2014; Lobb, 2014; Lobb and Jaeger, 2015; Murer et al., 1997). Such changed input patterns are likely to trigger homeostatic plasticity mechanisms in BGMT that can then further change the integrative properties of the thalamocortical pathway neurons and may lead to maladaptive activity patterns contributing to Parkinsonian motor dysfunction.

We addressed the question of BGMT changes in cellular properties after unilateral 6-OHDA treatment in mice that had undergone a minimum of 1 month of dopamine depletion allowing homeostatic compensation mechanisms to take place. We found a pronounced increase in excitability due to a decrease in M-type potassium current. This outcome will result in increased BGMT spiking activity that would be compensatory to increased levels of inhibitory BG input. However, the concomitant increase of T-type calcium channel elicited post-inhibitory rebound LTS bursting would also be likely to lead to an increased burstiness of BGMT output, particularly in the presence of synchronized inhibitory input bursts originating from the SNr. Increased motor thalamic LTS bursting has been found in MPTP treated primates (Devergnas et al., 2016; Magnin et al., 2000). While assessment of BGMT bursting in awake rodents is yet lacking, a recent study shows increased bursting following 6-OHDA treatment recorded under urethane anesthesia (Di Giovanni et al., 2020). Interestingly, in this study, an acute dopamine depletion lead to a decrease in BGMT firing rate and increase in thalamic GABA transmission, whereas the chronic 6-OHDA depleted state resulted not in a firing rate change, but in increased bursting, supporting the notion of an intervening homeostatic mechanism as dopamine depletion persists.

The M-type current (carried by KCNQ channels, more recently renamed as Kv7 channels) has been previously shown to be present in thalamocortical neurons in sensory thalamic neurons (Kasten et al., 2007), where a block by XE-991 moderately enhanced firing rate and lowered rheobase. More recently, both Kv7.2 and Kv7.3 were found abundantly expressed in the ventrobasal thalamus, and the number of LTS spikes following hyperpolarization increased with XE-991 block (Cerina et al., 2015). Our results are in good agreement with these previously observed effects of XE-991 in thalamocortical neurons, though they had not been previously demonstrated in motor thalamic areas. The complete lack of XE-991 effects following chronic 6-OHDA depletion we observed indicate a strong reduction of M-type current in this state. As supported by our modeling results, this effect can single-handedly account for the majority of our observations: A decrease in rheobase, an increase in spike rates, and an increase in LTS spikes following hyperpolarizing current injection. Interestingly, a homeostatic regulation of Kv7 channels has been previously observed in hippocampus and depends on L-type calcium channel signaling (Wu et al., 2008). Decreased BGMT spike rates following initial states of dopamine depletion would reduce L-type calcium currents and depotentiate M-type current according to this mechanism. KCNQ/Kv7 channels have also been localized to other key structures controlling brain rhythmic activity and neuronal synchronization including the SNr and the reticular nucleus of thalamus (Cooper et al., 2001). Thus, a more widespread function of these channels in controlling synchronization in basal ganglia circuits following dopamine depletion is quite possible.

The hyperpolarization activated current (IH) carried by HCN channels has been found to profoundly affect excitability and spiking regularity (Chan et al., 2004) as well as responses to synaptic inputs (Atherton et al., 2010) in a variety of cell types and it can undergo activity dependent regulation (Wang et al., 2002). This current is present in thalamus where it is known to regulate rhythmic activity patterns (Kanyshkova et al., 2009). Further, an HCN channelopathy has been found to be associated with dopamine depletion in globus pallidus neurons (Chan et al., 2011). When we tested BGMT neurons for changes in sag current following 6-OHDA treatment we identified a significant increase in sag (Fig. 3). While a detailed voltage-clamp analysis of HCN current was beyond the scope of our study, our modeling results suggest that the observed magnitude of sag increase cannot be accounted for by a decrease in M-current alone, suggesting an additional upregulation in IH in the 6-OHDA treated condition. Following findings for subthalamic neurons, such upregulation is likely to lead to a shunting of inhibitory inputs that would limit T-type Ca++ channel de-inactivation and reduce bursting (Atherton et al., 2010).

In conclusion, our finding of homeostatic changes in M-type potassium current gives a clear indication that cellular changes in BGMT could play a considerable role in the changes of thalamic activity in Parkinsonian conditions, and notably increased bursting and synchrony. Due to the traditional view of the BGMT as a mere relay of a BG rate code, changes in BGMT activity in behaving animals following dopamine depletion remain poorly studied to date. Recent studies from healthy rodents clearly indicate, however, that VM, which is the biggest component structure of BGMT, shows a closed excitatory loop with ALM cortical activity and indeed cortical persistent activity during motor preparation collapses when VM is inactivated (Guo et al., 2018; Guo et al., 2017). Additionally, VM neurons show a complex set of activity changes during sensory cued motor decision making tasks related to sensory cues, motor preparation, and motor execution (Catanese and Jaeger, 2020; Guo et al., 2017). Therefore any changes in VM excitability and synchrony are likely to play an important role in motor preparation.

## Acknowledgements

Edyta K Bichler current affilation: Department of Physiology, Emory University School of Medicine, Atlanta, GA 30322

The authors wish to acknowledge the considerable help provided by the anatomy core of the Emory Udall center, and specifically Dr. Adriana Galvan and Susan Jenkins. In addition, many thanks go to Dr. Su Li for supporting Matlab code development for data analysis. This work was supported by NINDS P50-NS098685 (Jaeger, Project 1 PI, Wichmann, Center PI) and NINDS 1R01NS111470 (Jaeger, PI).

## Notes

There is no conflict of interest for any of the authors.

### Competing Interest Statement

The authors have declared no competing interest.

